# CGRP inhibits SARS-CoV-2 infection of bronchial epithelial cells and its pulmonary levels correlate with viral clearance in critical COVID-19 patients

**DOI:** 10.1101/2024.01.05.574360

**Authors:** Caio César Barbosa Bomfim, Hugo Genin, Andréa Cottoignies-Callamarte, Sarah Gallois-Montbrun, Emilie Murigneux, Anette Sams, Arielle R Rosenberg, Sandrine Belouzard, Jean Dubuisson, Olivier Kosminder, Frédéric Pène, Benjamin Terrier, Morgane Bomsel, Yonatan Ganor

**Author notes:** **Corresponding author**: Yonatan Ganor.

## Abstract

Upon infection with severe acute respiratory syndrome coronavirus 2 (SARS-CoV-2), patients with critical coronavirus disease 2019 (COVID-19) present with life-threatening respiratory distress, pulmonary damage and cytokine storm. One unexplored hub in COVID-19 is the neuropeptide calcitonin gene-related peptide (CGRP), which is highly abundant in the airways and could converge in multiple aspects of COVID-19-related pulmonary pathophysiology. Whether CGRP affects SARS-CoV-2 infection directly remains elusive. We show that in critical COVID-19 patients, CGRP is increased in both plasma and lungs. Importantly, CGRP pulmonary levels are elevated in early SARS-CoV-2-positive patients, and restore to baseline upon subsequent viral clearance in SARS-CoV-2-negative patients. We further show that CGRP and its stable analogue SAX directly inhibit infection of bronchial Calu-3 epithelial cells with SARS-CoV-2 Omicron and Alpha variants in a dose-dependent manner. Both pre- and post-infection treatment with GRRP and/or SAX is enough to block SARS-CoV-2 productive infection of Calu3 cells. CGRP-mediated inhibition occurs via activation of the CGRP receptor and involves down-regulation of SARS-CoV-2 entry receptors at the surface of Calu-3 cells. Together, we propose that increased pulmonary CGRP mediates beneficial viral clearance in critical COVID-19 patients, by directly inhibiting SARS-CoV-2 infection. Hence, CGRP-based interventions could be harnessed for management of COVID-19.

**Brief summary:** Pulmonary levels of the neuropeptide CGRP are increased in critical COVID-19 patients, and could clear virus by directly inhibiting SRAS-CoV-2 infection of bronchial epithelia cells.

## Introduction

COVID-19 is characterized by pulmonary symptoms ranging from mild/moderate upper-airway disease to severe/critical acute respiratory distress syndrome (ARDS) and lung damage (1). Life threatening critical COVID-19 develops in patients with pre-disposing risk factors, including age of >65y, being male, and presenting with co-morbidities, such as hypertension, obesity and diabetes (2). These patients develop a cytokine storm characterized by elevated levels of pro-inflammatory cytokines such as interleukin 6 (IL-6) and tumor necrosis factor alpha (TNFα) (3) but impaired type I interferon responses (4–6), promoting altogether viral sepsis (7). Airway epithelial cells, which are key elements in the host innate immune response against respiratory infections (8), are a major cellular source for secretion of these pro-inflammatory cytokines (9, 10) upon SARS-CoV-2 sensing (11). In turn, SARS-CoV-2 replicates in epithelial cells throughout the airways (12), with a decreasing infection gradient from upper bronchial/bronchiolar to lower alveolar epithelial cells (13).

The 37 amino acids neuropeptide CGRP (14) is one unexplored hub in COVID-19. CGRP is highly abundant in the airways, where it is principally secreted by sensory nerve fibers (15) that innervate the lungs (16), and by rare bronchial epithelial cells with specialized sensory function termed pulmonary neuroendocrine cells (PNECs) (17). Due to its immunodulatory and vasodilatory functions, CGRP might intersect with multiple COVID-19 pulmonary symptoms, which could result in opposite consequences. Accordingly, CGRP is a key regulator of inflammatory processes and affects the release of pro-inflammatory cytokines. While CGRP increases IL-6 secretion from bronchial epithelial cells (18), it also decreases TNFα levels in murine models of bacterial sepsis in which CGRP levels are elevated (19). In addition, CGRP is a potent vasodilator (20) believed to be secreted during local hypoxia, with beneficial actions in cardiovascular diseases including hypertension (21, 22). Hence, CGRP levels/effects vary in pathologies that characterize and/or represent risk factors for development of critical COVID-19. CGRP further promotes lung tissue repair and regeneration (23), by inducing bronchial epithelial cells migration (24) and wound healing (25). These studies indicate that CGRP plays both harmful and protective roles in airways physiopathology.

Besides its immunomodulatory and vasodilatory roles, CGRP also exerts anti-viral functions, as we previously discovered (26–30). Indeed, CGRP strongly inhibits infection of antigen-presenting Langerhans cells (LCs) with human immunodeficiency virus type 1 (HIV-1) and herpes simplex virus types 1 and 2 (HSV-1/2) (26–30). One shared CGRP-mediated anti-viral mechanism involves modulation of surface expression of different viral attachment and entry receptors. In particular, CGRP: i) enhances surface expression of atypical double-trimmers of the LC-specific pathogen-recognition lectin langerin, i.e. the receptor of both HIV-1/HSV-2, resulting in reduced viral capture (30); ii) decreases surface expression of the HIV-1 entry co-receptor CCR5 (27), and the HSV-1 entry receptor 3-O sulfated heparan sulfate (30).

Herein, we explored the interplay between CGRP and COVID-19 *in-vivo* and investigated whether CGRP exerts direct anti-SARS-CoV-2 effects when acting on pulmonary epithelial cells.

## Results

### CGRP levels are increased in SARS-CoV-2-positive critical COVID-19 patients

To explore the involvement of CGRP in COVID-19, we used some of our previously described plasma samples (4), obtained from COVID-19 patients with different disease severities, namely mild/moderate, severe and critical. Control samples included plasma from patients with non-COVID-19-related pathologies as we described (31), as well as from otherwise healthy individuals at time of sampling that we also described (32). As shown in Table 1, all cases had mean age of >65y, were mostly males, and were matched for both age and sex. When compared to all other groups, critical COVID-19 patients had significantly higher incidence of smokers (69%, p=0.0473). These patients also had higher incidence of hypertension, yet not reaching statistical significance (62%, p=0.0790). On multivariate analysis in this cohort, smoking was associated with critical COVID-19 (odds ratio (OR) 6.8 [95% confidence intervals (CIs) 1.3-42.8]; p=0.0259). We measured CGRP in these plasma samples by an enzyme immunoassay (EIA), and found that critical COVID-19 patients had significantly elevated CGRP levels, when compared to the other groups of COVID-19 patients, as well as to non-COVID-19 patients and healthy controls (Figure 1A).

**Figure 1.**
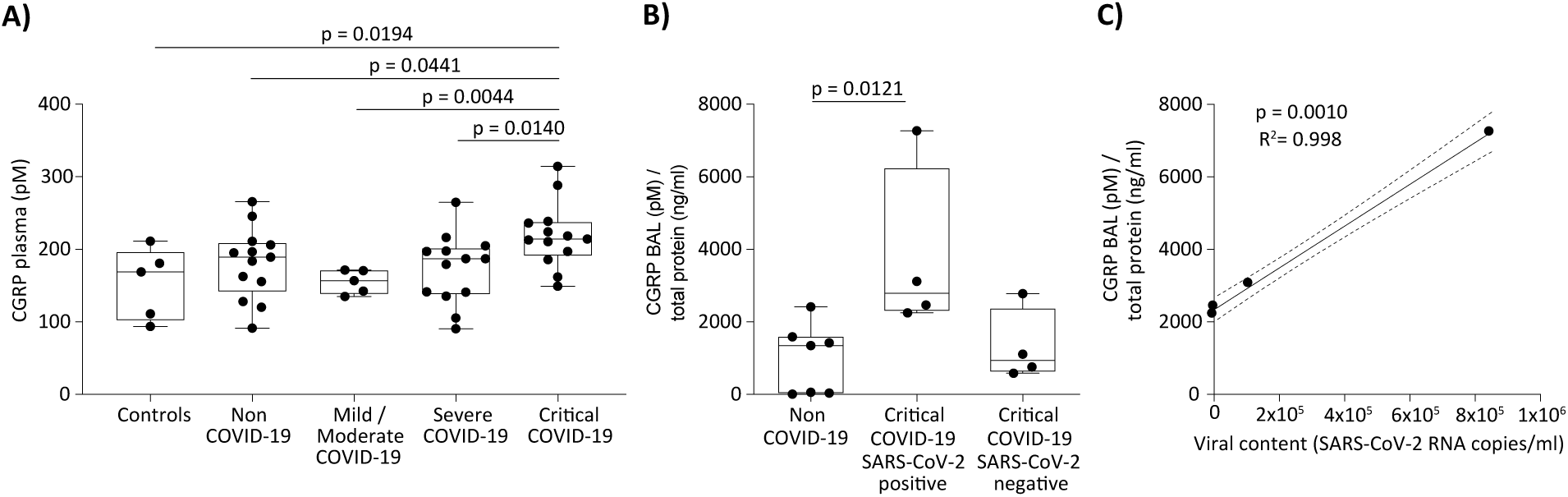
CGRP levels are increased in critical COVID-19 patients. **(A, B)** CGRP levels were determined using an EIA, in plasma of COVID-19 patients with different disease severities, non-COVID-19 patients and healthy controls (A), and in BALs of SARS-CoV-2-positive or negative critical COVID-19 and non-COVID-19 patients (B). In (B), CGRP levels were normalized to that of BAL total protein content. Shown are box and whisker plots with exact p values calculated with the Mann-Whitney test. **(C)** Correlation between SARS-CoV-2 viral content expressed as RNA copies / ml and normalized CGRP levels, in BALs of SARS-CoV-2-positive critical COVID-19 patients. Solid line denotes a linear correlation with numbers representing the corresponding p and R^2^ values, and broken lines denote 95% CIs.

**Table 1.**
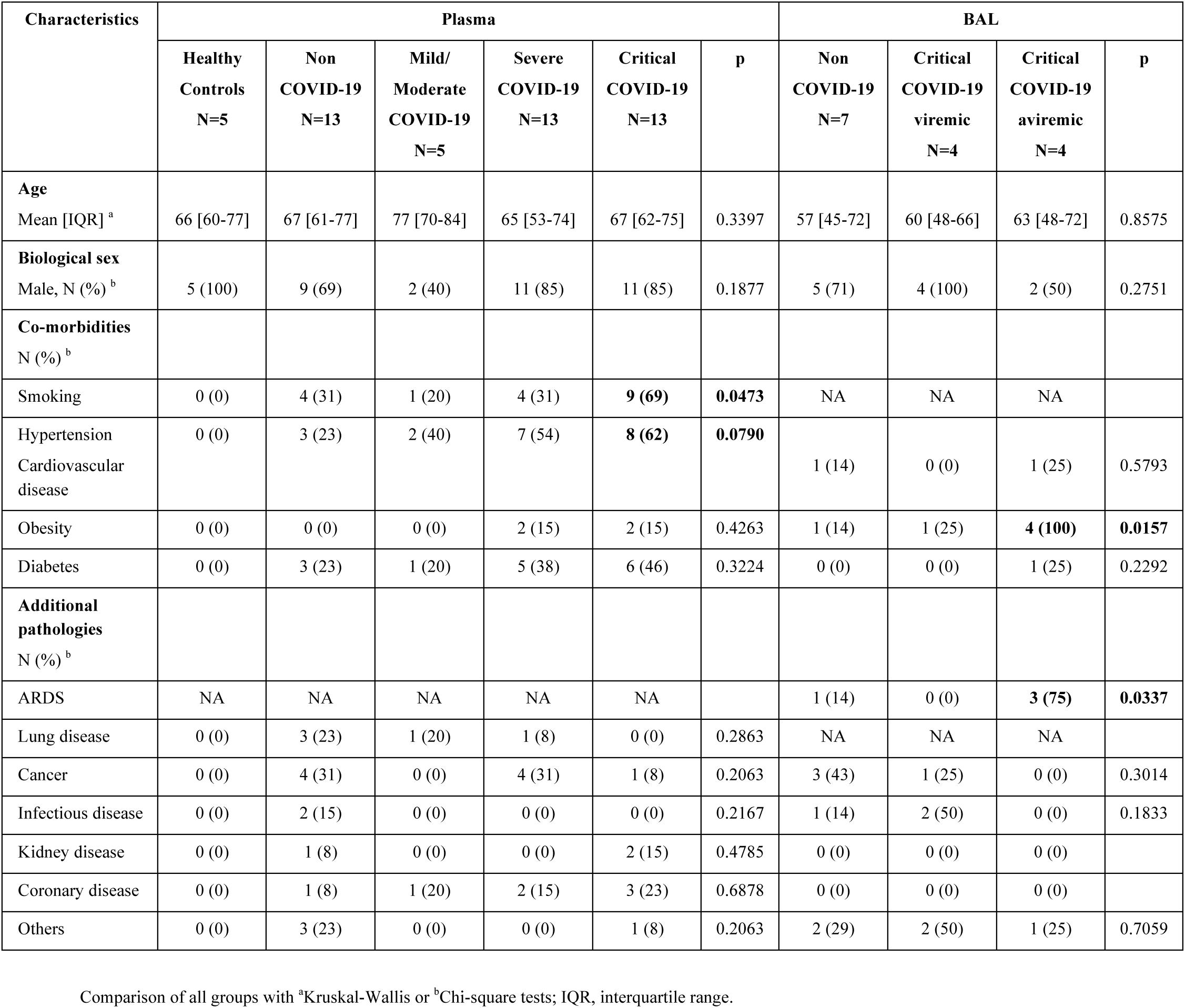
Clinical characteristics of the study patients.

SARS-CoV-2 detection and isolation from non-respiratory specimens, including blood, are unsuccessful in most studies (33). Therefore, we also measured CGRP in bronco-alveolar lavages (BALs) of additional critical COVID-19 and non-COVID-19 patients (i.e. serving as controls), which were included in our other study (34). We previously quantified the viral content in these BAL samples and reported that SARS-CoV-2 presence in BALs (i.e. SARS-CoV-2-positive patients) corresponds to an early phase of viral replication, whereas SARS-CoV-2 absence in BALs (i.e. SARS-CoV-2-negative patients) corresponds to a later phase of the infection when virus has been cleared from the lungs (34). These cases were matched in terms of age and sex (see table 1), and were not statistically different from the cases from which we obtained plasma samples in terms of both age (p=0.2477) and sex (p=0.2197). When compared to the SARS-CoV-2-positive critical COVID-19 and non-COVID-19 patients, SARS-CoV-2-negative critical COVID-19 patients had significantly higher incidence of obesity (100%, p=0.0157) and ARDS (75%, p=0.0337). We measured CGRP by EIA in these BAL samples as above, and normalized CGRP levels to that of total BAL protein (i.e. to account for potential differences in protein content upon BALs collection). When compared to the non-COVID-19 patients, normalized CGRP levels were significantly higher in BALs of SARS-CoV-2-positive critical COVID-19 patients (Figure 1B). Moreover, these levels significantly and positively correlated with SARS-CoV-2 viral content in these samples (Figure 1C). Interestingly, in BALs obtained from SARS-CoV-2-negative critical COVID-19 patients that correspond to a late stage of the disease, normalized CGRP levels were restored to baseline and were not different from that in the control group (Figure 1B). These results show that CGRP levels increase in critical COVID-19 patients and correlate with their pulmonary viral content.

### CGRP and its analogue SAX inhibit SARS-CoV-2 infection of Calu-3 cells

SARS-CoV-2 content in BALs (i.e. reflecting pulmonary viral content) is transitory, suggesting that CGRP might interfere with viral replication. Thus, we next explored the direct effects of CGRP on SARS-CoV-2 infection, using the well-characterized bronchial epithelial cell line Calu-3 (35) that is permissive to infection with different SARS-CoV-2 variants (36). We first confirmed that Calu-3 cells were productively infected, by pulsing confluent Calu-3 monolayers with SARS-CoV-2 for 4h, alone or following pre-incubation with the antiviral drug remdesivir (RDV). Four days later, viral replication was evaluated by intracellular cell expression of the SARS-CoV-2 spike protein using flow cytometry. These experiments showed that Calu-3 cells were infected with both Omicron (Figure 2A) and Alpha (Figure 2B) variants. As expected, infection was productive and was completely inhibited by RDV for both SARS-CoV-2 variants (Figure 2A, B).

**Figure 2.**
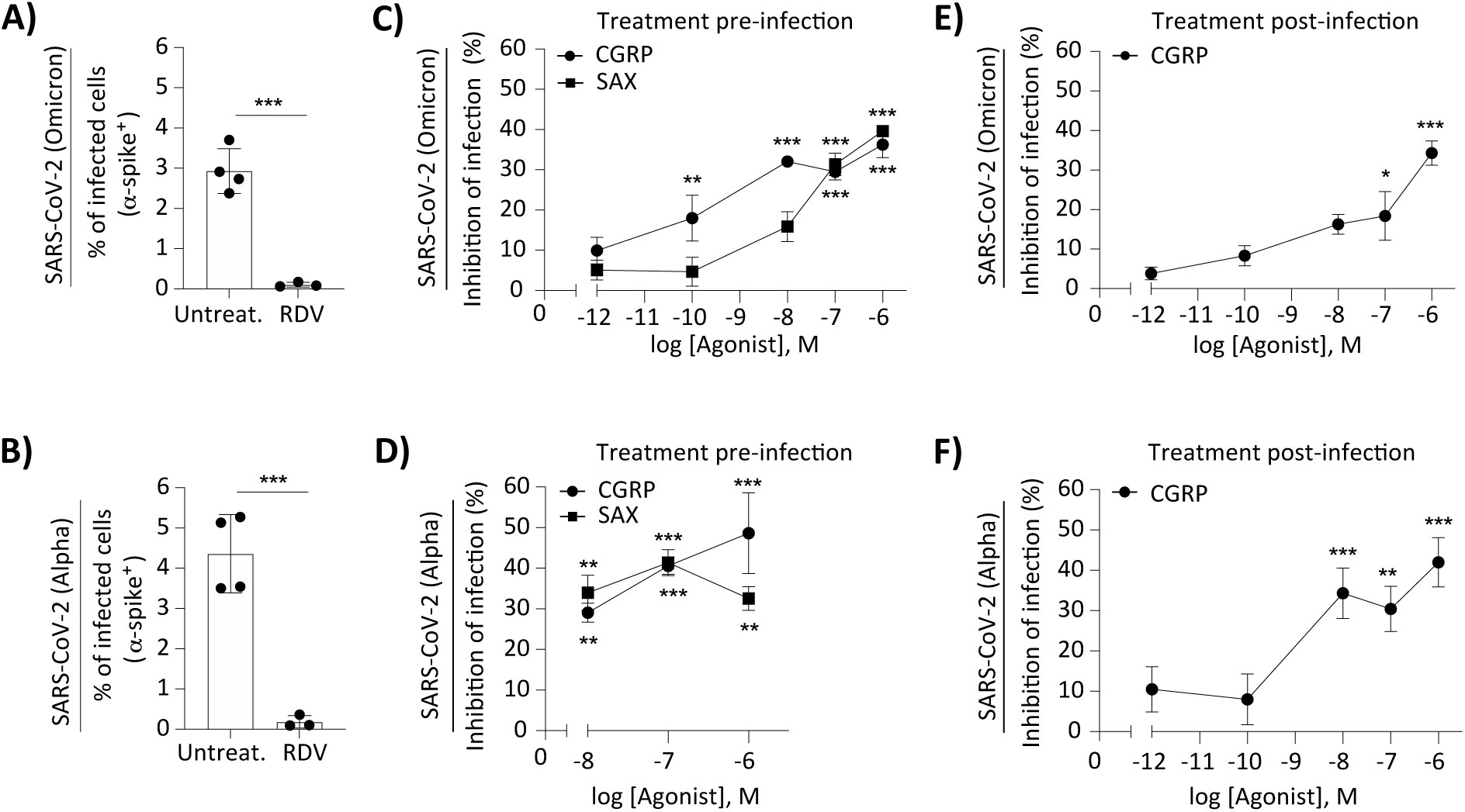
CGRP and SAX inhibit SARS-CoV-2 infection of Calu-3 cells. **(A, B)** Calu-3 cells were left untreated or pre-incubated for 24h at 37°C with RDV, washed, pulsed with the SARS-CoV-2 Omicron (A) or Alpha (B) variants for 4h at 37°C, washed again, and further incubated for 4 days. Infection was then determined by intracellular staining using an anti-spike Ab and flow cytometry. Graphs represent mean±SEM (n=4 independent experiments) percentages of spike-positive infected cells. **(C, D)** Calu-3 cells were left untreated or pre-treated for 24h at 37°C with the indicated molar concentrations of CGRP or SAX. The cells were then washed, pulsed with SARS-CoV2 Omicron (C) or Alpha (D) variants and examined for infection by flow cytometry as above. Graphs represent mean±SEM (n=3 independent experiments) percentages of SARS-CoV2 infection inhibition, normalized against untreated cells serving as the 100% set point (i.e. no inhibition). **(E, F)** Untreated Calu-3 cells were pulsed with SARS-CoV2 variants as above, washed, and post-treated with the indicated molar concentrations of CGRP that was kept throughout the 4 days of culture. Results (n=3 independent experiments) are shown as in (C, D). In all experiments, statistical significance was evaluated by the Student’s t-test.

Next, Calu-3 cells were pre-treated for 24h at 37°C with CGRP within its concentration range that inhibits LCs-mediated HIV-1 and HSV-1/2 infection, as we reported (29, 30). The cells were also pre-treated with the CGRP metabolically stable analogue SAX (serinyl-CGRP_2-37_- amide with an albumin binding fatty acid moiety in the N-terminus) that, like CGRP, exerts protective cardiovascular effects (37–40) and anti-viral functions (29, 30), as we reported. The cells were next washed, pulsed with SARS-CoV-2 variants for 4h, and infection was determined by flow cytometry as above. CGRP and SAX significantly inhibited infection of Calu-3 cells with both Omicron (Figure 2C) and Alpha (Figure 2D) variants, in a dose-dependent manner. The potency of SAX was lower than that of CGRP for the Omicron variant (Figure 2C; p=0.0268), which is in accordance with our previous studies (37, 39).

Calu-3 cells were also pre-treated for 24h at 37°C with 10^-6^M CGRP (i.e. its highest effective concentration we tested), pulsed with the SARS-CoV-2 Alpha variant for 4h, and viral production was evaluated 24h later by quantifying viral genome copies in the culture supernatants. CGRP reduced viral production from Calu-3 cells, with mean±SEM (derived of n=4 different experiments) relative SARS-CoV-2 RNA folds of 1.0±0.13 for untreated cells vs. 0.61±0.08 for CGRP-treated cells (p=0.0432, Student’s t-test).

To further evaluate the mechanisms of action of CGRP, Calu-3 cells were first pulsed with the SARS-CoV-2 variants, washed, and CGRP was added after the viral pulse and maintained throughout the 4 days culture period. Such CGRP post-treatment significantly inhibited infection of Calu-3 cells with both SARS-CoV-2 variants, in a dose-dependent manner (Figure 2E, F). Moreover, the extent of inhibition mediated by CGRP pre- (Figure 2C, D) and post-infection (Figure 2E, F) was similar. These results indicate that CGRP directly inhibits SARS-CoV-2 replication in bronchial epithelial cells.

### CGRP-mediated inhibition of SARS-CoV-2 infection involves activation of the CGRP receptor in Calu-3 cells

The heteromeric CGRP receptor complex is composed of calcitonin receptor-like receptor (CLR) and receptor activity modifying protein 1 (RAMP1; important for surface trafficking of the receptor complex), as well as the intracellular receptor component protein (RCP; important for signaling of this receptor) (41). Primary airway epithelial cells were reported to express the CGRP receptor (42), but data is missing for Calu-3 cells. Hence, we stained Calu-3 cells with commercially available antibodies (Abs) that are validated for flow cytometry and directed against extracellular epitopes of human CLR and RAMP1. Approximately 10% of Calu-3 cells co-expressed CLR and RAMP1 on their surface (Figure 3A), indicating the presence of a potentially functional CGRP receptor.

**Figure 3.**
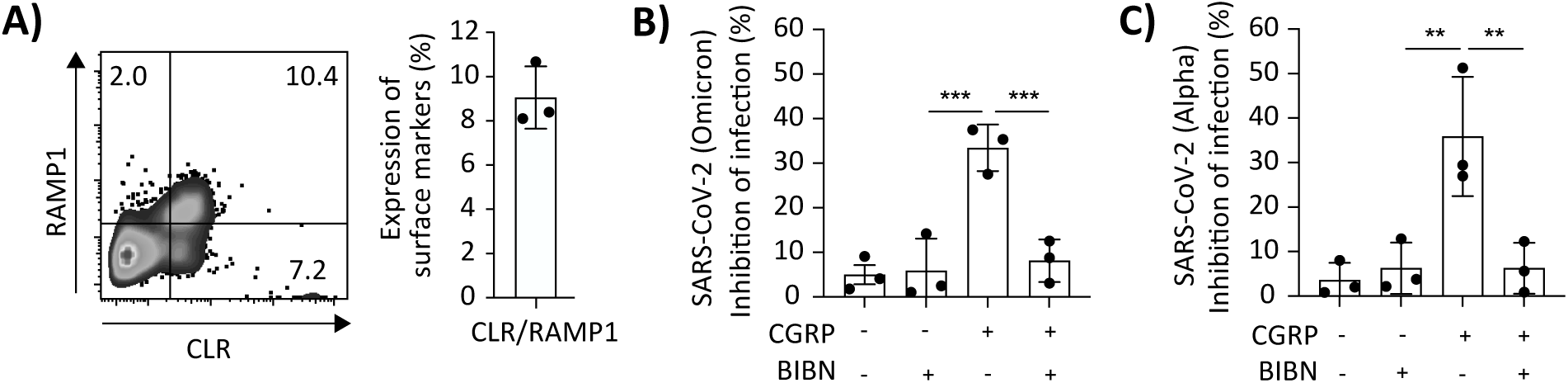
CGRP inhibits SARS-CoV-2 infection of Calu-3 cells by activating its receptor. **(A)** Representative flow cytometry dot plot showing surface expression of CLR and RAMP1 in Calu-3 cells, with graph showing mean±SEM (n=3 independent experiments) percentages of double positive cells. **(B, C)** Calu-3 cells were left untreated or pre-treated for 24h at 37°C with 10^-6^M CGRP. When indicated, the CGRP receptor antagonist BIBN was added at 10^-6^M 15min before addition of CGRP. Cells were then washed, pulsed with the SARS-CoV-2 Omicron (B) or Alpha (C) variants for 4h at 37°C, washed again, further incubated for 4 days, and infection was determined by flow cytometry. Graphs represent mean±SEM (n=3 independent experiments) percentages of SARS-CoV2 infection inhibition, and statistical significance was evaluated by the Student’s t-test.

To determine the implication of the CGRP receptor in the anti-viral activity of CGRP, Calu-3 cells were first incubated for 15min with the CGRP receptor antagonist BIBN4096 (BIBN), followed by pre-treatment with 10^-6^M CGRP for 24h. The cells were then washed, pulsed with SARS-CoV-2 for 4h, and infection was evaluated 4 days later by flow cytometry as above. BIBN alone had no significant effect, but completely abrogated CGRP-mediated inhibition of infection with both Omicron (Figure 3B) and Alpha (Figure 3C) variants.

These results indicate that the anti-SARS-CoV-2 activity of CGRP is mediated via activation of its cognate receptor expressed at the surface of Calu-3 cells.

### CGRP reduces expression of SARS-CoV-2 entry receptors in Calu-3 cells

We next confirmed by flow cytometry the expression SARS-CoV-2 entry receptors at the surface of Calu-3 cells (Figure 4A), namely angiotensin-converting enzyme 2 (ACE2) and transmembrane protease serine 2 (TMPRSS2) (43). We then investigated whether CGRP affects the surface levels of these receptors, like its capacity to modulate expression of HIV-1 and HSV-1/2 entry receptors mentioned above. Calu-3 cells were left untreated or treated with 10^-6^M CGRP for 24h, and surface expression of ACE2 and TMPRSS2 was determined immediately after the 24h CGRP treatment or following washing and incubation for additional 24h. These experiments showed that surface expression of both ACE2 (Figure 4B) and TMPRSS2 (Figure 4C) was significantly decreased by approximately half following CGRP treatment, and was resorted the next day.

**Figure 4.**
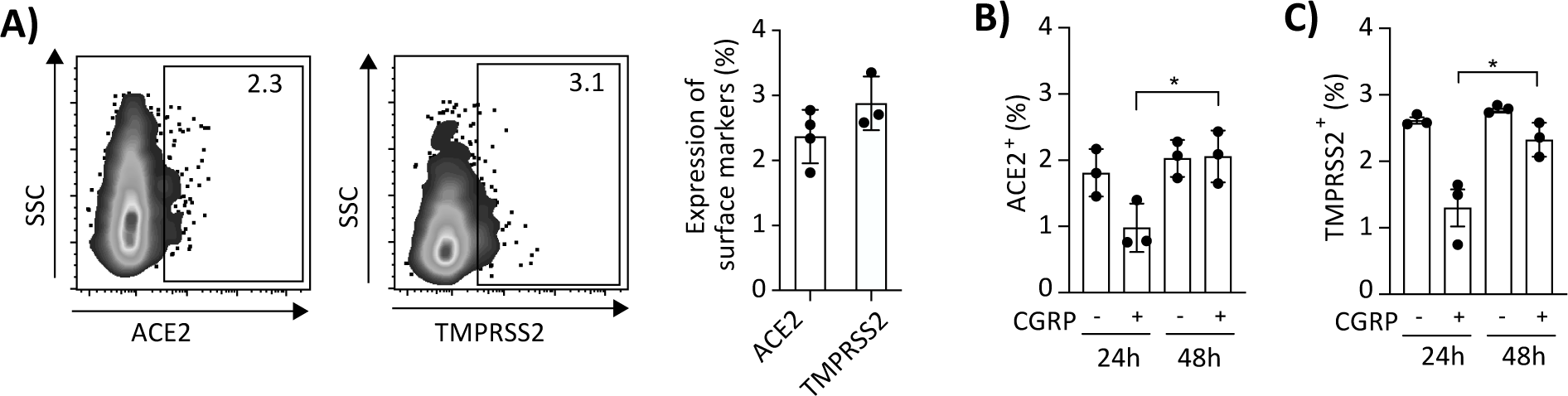
CGRP decreases ACE2 and TMPRSS2 surface expression in Calu-3 cells. **(A)** Representative flow cytometry dot plots showing surface expression of ACE2 or TMPRSS2 in untreated Calu-3 cells, with graphs showing mean±SEM (n=3-4 independent experiments) percentages of positive cells. **(B, C)** Calu-3 cells were left untreated or treated for 24h at 37°C with 10^-6^M CGRP, washed, and incubated for additional 24h. Surface expression of ACE2 and TMPRSS2 was determined immediately after CGRP treatment (t=24h) or following washing and further incubation (t=48h). Graphs represent mean±SEM (n=3 independent experiments) percentages of positive cells, and statistical significance was evaluated by the Student’s t-test.

These results suggest that CGRP inhibits Calu-3 infection by limiting SARS-CoV-2 entry in Calu-3 cells.

## Discussion

The rapid development of vaccines against SARS-CoV-2 and direct-acting antivirals was crucial for management of the COVID-19 pandemic. Although theses clinical interventions prevent severe disease, vaccine-induced protection weakens in months and SARS-CoV-2 variants escape vaccine-acquired immunity (2). As these strategies fail to fully block viral transmission, identifying endogenous pulmonary factors limiting SARS-CoV-2 infection could offer novel preventive and therapeutic approaches complementing vaccination.

We show that CGRP levels are increased in the plasma of critical COVID-19 patients. Moreover, we tested for the first time CGRP levels in BALs of COVID-19 patients, namely samples representing the tissue site of SARS-CoV-2 replication, and observed similar CGRP elevation. Importantly, CGRP increases only when BALs from critical COVID-19 patients contain SARS-CoV-2. This suggests that CGRP increases when virus replicates in the lungs of these patients, helping to control pulmonary SARS-CoV-2 replication. As plasma CGRP is considered as ‘spillover’ from its tissue production sites, the changes in CGRP levels we observed in the plasma of critical COVID-19 patients correspond to different stages of their disease. Future studies should now quantify CGRP levels in BALs of non-critical COVID-19 patients, which we speculate will not contain elevated CGRP.

Interestingly, the overall outcome we observe is elevation of CGRP in both plasma and SARS-CoV-2-positive BALs, although the samples tested herein were obtained from critical COVID-19 patients with several co-morbidities that have opposite effects on CGRP. Specifically, our samples were obtained from patients with high incidence of smoking/hypertension (for plasma) or obesity (for SARS-CoV-2-negative BALs), which associate with either increased (i.e. smoking (44) and obesity (22)) or decreased (i.e. hypertension (21)) CGRP levels. When the plasma samples of critical COVID-19 patients were stratified according to smoking status, CGPR levels were higher in smokers compared to non-smokers (i.e. mean±SEM of 229.3.0±16.6 vs. 197.0±12.0 respectively), yet not reaching statistical significance (p=0.1986, Mann-Whitney test). These results suggest that the net effect on CGRP levels cannot be attributed to a single clinical parameter.

Of note, age and sex are additional factors that should be considered, as plasma CGRP levels decrease with age (22) and are lower in males (45, 46). These observations could explain why a previous study concluded that plasma CGRP levels were lower in COVID-19 patients, as the control group in that study included healthy volunteers that were significantly younger than the COVID-19 patients tested. Nevertheless, although not reaching statistical significance, CGRP levels were higher in COVID-19 patients admitted to the intensive care unit (ICU) compared to patients at the general ward or asymptomatic (47). Similarly, a second study reported that plasma CGRP levels were higher in a cohort of COVID-19 patients that excluded ICU patients. Yet, cases were younger and included males and females at equal proportions (48), differing from our cohort that includes principally elderly males. Finally, a third study evaluated plasma CGRP in a cohort of moderate to severe COVID-19 patients, but without comparison to any controls. This study included elderly and mostly male patients, and reported that CGRP levels were higher in patients with negative disease evolution, including eventual ICU admission (49). Collectively, and together with our results described herein, we conclude that elevated CGRP is a hallmark of early COVID-19.

The control of SARS-CoV-2 replication by CGRP, we observed *in-vitro*, suggests the induction of beneficial CGRP-mediated protective mechanisms. Indeed, in other lung pathologies such as asthma and cystic fibrosis, elevated CGRP is not causing damage but is rather a compensatory protective response to ameliorate the damage (50). Similarly, increase in plasma CGRP levels in sepsis is considered a compensatory mechanism by which septic shock is attenuated (51). Our results indicate that this compensatory response probably also occurs in COVID-19, as we find that CGRP directly inhibits SARS-CoV-2 replication in bronchial epithelial cells. Such inhibition involves activation of the CGRP receptor and leads to down-regulation of SARS-CoV-2 entry receptors. Importantly, reduced infection following CGRP pre-treatment correlates with the decrease in ACE2/TMPRSS2 surface expression at the time of viral pulse. In CGRP post-treatment, we speculate that the maintenance of CGRP throughout the infection period provides a sufficient time frame, permitting CGRP to exert a similar ACE2/TMPRSS2 decreasing effect, which is similarly accompanied by reduced infection. CGRP post-treatment could also impair viral propagation by reducing ACE2/TMPRSS2 surface expression on cells not yet infected after SARS-CoV-2 pulse. We therefore propose that future CGRP-based formulations might be useful for COVID-19 prevention. Such formulations might also be useful as COVID-19 treatments, at an early time point when the virus is still present in the airways. Our results also extend the anti-viral function of CGRP to pulmonary viruses and cells, in addition to that exerted by acting on immune cells that are targeted by sexually transmitted viruses, as we previously discovered.

Different cell types secrete CGRP in the airways, principally sensory nerves and PNECs but also non-neuronal cells, including macrophages (52) and epithelial cells (42, 53). Previous studies reported that SARS-CoV-2 preferentially targets ciliated and secretory lung epithelial cells (54). These results are in line with the SARS-CoV-2 permissiveness of Calu-3 cells, which give rise to both cell types upon their differentiation in submerged and air-liquid culture conditions (55). Interestingly, while bronchial epithelial cells secrete pro-inflammatory cytokines upon sensing of SARS-CoV-2, we did not observe induction of CGRP secretion from SARS-CoV-2-infected Calu-3 cells (data not shown). Hence, it would be instrumental to determine precisely CGRP cellular sources during SARS-CoV-2 infection. Collectively, we reveal a novel anti-viral function of CGRP and suggest that its direct anti-SRAS-CoV-2 function mediates beneficial pulmonary viral clearance in critical COVID-19. We further argue that caution should be taken when considering the use of CGRP receptor antagonists for COVID-19 management (56), which would counteract these direct protective effects of CGRP against SRAS-CoV-2 infection.

## Methods

### Plasma and BAL levels of CGRP

Plasma samples were obtained from different groups of COVID-19 patients, which were hospitalized in either the internal medicine unit (IMU) or ICU at the Cochin Hospital (Paris). These patients were described in our previous study (4) and presented with different COVID-19 disease severities at sampling date. In the current study, we used samples from n=5 mild/moderate patients at the IMU, n=13 severe patients at the IMU and n=13 critical patients at the ICU. As control, we used plasma samples from n=13 patients admitted due to various non-COVID-19-related pathologies that we described in another previous study (31), as well as from n=5 otherwise healthy individuals that we recently described (32).

BAL samples were obtained from additional critical COVID-19 patients hospitalized at the ICU of the Cochin (Paris) or Ambroise Paré (Boulogne-Billancourt) Hospitals, which we previously described (34). Herein, we included n=4 SARS-CoV-2-positive and n=4 SARS-CoV-2-negative critical COVID-19 patients, as well as n=7 patients admitted due to non-COVID-19-related pathologies.

Plasma and BAL CGRP levels were determined using a competitive CGRP EIA kit (Phoenix Pharmaceuticals), and total BAL protein content was determined using the Pierce bicinchoninic acid assay (BCA) kit (Thermo Scientific), both according to the manufacturers’ instructions.

### Cells and viruses

The human bronchial Calu-3 epithelial cell line was obtained from American type Culture Collection (ATCC). Cells were cultured in Dulbecco’s Modified Eagle’s Medium (DMEM; Gibco) supplemented with 10% fetal calf serum (FCS; Eurobio), 2mM L-glutamine, 1% non-essential amino acids, 100U/mL penicillin, and 100µg/mL streptomycin (Gibco). Cells were incubated at 37°C and 5% CO_2_ and were detached using 0.05% trypsin (Gibco). SARS-CoV-2 included the Alpha and Omicron variants we described previously (57), and viral stocks were prepared and titrated using Vero-E6 cells as before (58).

### Calu-3 infection

Calu-3 cells were plated in 48-wells plate (2.5X10^5^ cells/well), incubated overnight at 37°C, and pre-treated for 24h with the indicated molar concentrations of CGRP (Sigma) or its stable analogue SAX (40), as we previously described (29, 39). When indicated, the CGRP receptor antagonist BIBN4096 (Sigma) was added 15min before addition of agonists, and the antiviral drug RDV (MedChemExpress) was added at 10μM overnight before infection. Cells were next washed, pulsed for 4h with SARS-CoV-2 variants at multiplicities of infection of 0.05 (for Alpha) or 0.0015 (for Omicron), and further incubated for 4 days. In some experiments, CGRP was added after the pulse period and kept until the end of the culture. Next, cells were detached with trypsin, transferred to 96 round-bottom wells plate, and fixed/permeabilized with the Cytofix/Cytoperm kit (BD Bioscience). Cells were then stained for 30min on ice with Alexa-Fluor-488-conjugated mouse monoclonal antibody (mAb) directed against the SARS-CoV-2 spike protein (R&D systems; clone #1035206, diluted 1:50 in Perm/Wash buffer (BD)). Infection was determined by flow cytometry, using a Guava easyCyte Flow Cytometer (Merck-Millipore) and analyzed by FlowJo software (FlowJo LLC).

In some experiments with the Alpha variant, the culture supernatants were collected 24h following the viral pulse, and infectious viral particle production was quantified by reverse transcription (RT)-qPCR, as we previously described (59).

### Surface expression of SARS-CoV-2 entry receptors

Calu-3 cells (5×10^4^ cells/well, in 96 round-bottom wells plate) were surface stained with fluorescein-isothiocyanate-conjugated rabbit-anti-human CLR (Alomone labs; 1:100). Cells were also surface stained with unconjugated rabbit-anti-human RAMP1 (Proteintech; 1:100), ACE2 (Biorbyt; 1:25) or TMPRSS2 (Invitrogen; 1:10), followed by cyanine-5-conjugated donkey-anti-rabbit-IgG secondary Ab (Jackson Immunoresearch; 1:200). All Abs were diluted in phosphate-buffered saline (PBS), and each staining step was performed with 30µl/well for 30min on ice. Cells were analyzed by flow cytometry, as above

### Statistical Analyses

Statistical significance was analyzed using Prism 9 software (GraphPad). For clinical data of COVID-19 patients and controls (Table 1), the non-parametric Kruskal-Wallis or Chi-square tests were used to compare continuous or categorical parameters across groups, respectively. A multivariate logistic regression model was applied to calculate ORs and 95% CIs, and the non-parametric unpaired Mann-Whitney test was used for pairwise comparisons of CGRP levels. Liner correlation was calculated for association between viral content and CGRP levels in BAL samples. For experimental data using Calu-3 cells, the Student’s t-test was used. CGRP and SAX dose-response curves were compared using the [log(inhibitor) vs. normalized response - variable slope] non-linear regression model. All tests were two-sided, and differences were considered significant when the p values were <0.0500, with asterisks denoting the degree of significance (*p<0.0500; **p<0.0050 and ***p<0.0005).

## Notes

**Conflict of interest statement**: The authors have declared that no conflict of interest exists.

### Competing Interest Statement

The authors have declared no competing interest.

